# Early visual cortex is not functionally necessary for temporal attention: A TMS study

**DOI:** 10.64898/2026.11.05.764720

**Authors:** Hsing-Hao Lee, Marisa Carrasco

## Abstract

Temporal attention enhances the selective processing of information at a specific point in time and improves perception. Previous neuroimaging studies investigating its neural mechanisms have either conflated temporal attention with temporal expectation or provided correlational evidence. Thus, whether early visual cortex is functionally necessary for temporal attention remains unknown. Here, we combined psychophysics with transcranial magnetic stimulation (TMS) to examine whether V1/V2 are functionally necessary and play a causal role in temporal attention. Observers performed an orientation-discrimination task in which two consecutive stimuli, T1 and T2, were presented in the periphery, preceded by a precue providing either valid or neutral temporal information. Double-pulse TMS was delivered under four conditions: at T1 (T1-TMS), at T2 (T2-TMS), at both T1 and T2 (both-TMS), or not at all (none-TMS), to assess its effects on temporal attention and the efficacy of the stimulation was verified by pupil size analysis. Behavioral results revealed robust temporal attention benefits on *d′*, reaction time, and the Balanced Integration Score (BIS) across all four TMS conditions, indicating that the benefits of temporal attention were unaffected by occipital TMS. Furthermore, pupil dilation was largest in the both-TMS condition, followed by the T1-TMS and T2-TMS conditions, and smallest in the none-TMS condition, providing physiological verification of the stimulation protocol. Together, these findings reveal the first causal evidence that the early visual cortex is not functionally necessary for endogenous temporal attention.

**Significance:** Temporal attention prioritizes sensory information at specific moments, yet its underlying neural mechanisms remain poorly understood due to a lack of direct causal evidence. Neuroimaging studies have implicated multiple areas, including early visual and higher-order cortical regions. Here, using transcranial magnetic stimulation (TMS) to disrupt neural activity in the early visual cortex, we found that behavioral benefits of temporal attention remained intact. These findings reveal that the early visual cortex is not functionally necessary for temporal attention. Critically, this study provides the first causal evidence that selectively and voluntarily attending to specific time points operates independently of early visual areas, suggesting that higher-order regions–such as the fronto-cingulate network–play a primary role.

## Introduction

Given the high energy cost of cortical computation and limited metabolic resources, the brain cannot process all visual information in the environment simultaneously. Selective visual attention is necessary to allocate limited resources according to task demands [1-3]. Visual attention can be deployed covertly across space [review, 2] or time [review, 4, 5], enhancing processing for attended locations or timepoints, while impairing unattended ones via a push-pull mechanism. More is known about the behavioral effects and neural bases of spatial attention than temporal attention. Here, we investigated the functional necessity of early visual cortex for temporal attention.

Top-down orientation of selective processing in time can be guided by both temporal attention (e.g., [6-12]) or temporal expectation (e.g., [13-16]). Temporal attention—the ability to prioritize and selectively process the specific time points based on behavioral relevance regardless of its predictability—and temporal expectation—the ability to formulate predictions about when a visual event is likely to occur—are fundamental cognitive processes that act together to distribute limited resources within a time window. These processes are closely related [17], as attention can be guided by expectation [18], but can be and should be disentangled [6, 7, 17]. It has been stated that converging evidence confirms that temporal attention enhances neural responses in early visual cortex and improves behavioral performance [19-24, review, 25]; however, this conclusion is premature. It remains unclear whether temporal attention and temporal expectation map neatly onto computationally or neurophysiologically dissociable brain mechanisms [26], as many previous functional magnetic resonance imaging (fMRI), electroencephalogram (EEG) and MEG studies conflate temporal attention and expectation.

Temporal expectation—typically manipulated by whether the target is presented at the predicted time point—engages early visual cortex by enhancing visually evoked responses [27, 28], reducing prestimulus occipital alpha power [29, 30], and modulating visual cortical activity [31-33]. Studies on temporal attention that did not isolate expectation have likewise reported enhanced early visual cortical activity in human and non-human primates when attention is directed toward an expected time point [21, 34, 35]. Together, these findings implicate early visual cortex in temporal expectation and its effects on visual processing, but see [36].

Recent magnetoencephalography (MEG) studies that manipulated temporal attention while keeping expectation constant provide evidence that early visual cortex plays a role in temporal attention. Although these studies observed no differences in visually-evoked responses between attended and unattended time points, temporal attention increased intertrial phase coherence in visual cortex and induced a phase shift toward the attended target [37]–suggesting that it adjusts the timing and reliability of visual cortical responses to prioritize information at attended moments. Additionally, temporal attention enhanced orientation-decoding accuracy in the left fronto-cingulate cortex, further suggesting that temporal attention sharpens sensory representations [38]. Furthermore, temporal attention enhanced average closeness centrality within the occipital lobe along a left-lateralized pathway, indicating increased sharing of target information along two pathways, occipito-fronto-cingulate and occipitotemporal. This suggests that visual cortex supports temporal attention by routing attended information to higher cortical regions [39].

However, whether early visual cortex is functionally necessary for temporal attention remains unknown because non-invasive neuroimaging techniques provide correlational evidence without manipulating neural activity. TMS overcomes this limitation by briefly disrupting local excitation and inhibition to establish the functional necessity of specific brain regions for perception and cognition [40, 41]. Critically, TMS effects are state dependent: suppressing active neuronal populations while disinhibiting less active ones. By manipulating brain state via psychophysical protocols—such as adaptation [42-45], covert attention [46, 47], or presaccadic attention [48] (for reviews [49-51])—researchers can predict TMS impacts behavioral performance on various tasks, such as orientation and motion [52].

To investigate whether early visual cortex is functionally necessary for temporal attention, we conducted a TMS-psychophysical experiment adapted from protocols used in covert spatial attention studies. Those spatial attention studies revealed that early visual cortex (V1/V2) is functionally necessary for involuntary (exogenous) attention [53, 54] but not voluntary (endogenous) attention [55], whereas the human homologue of the right frontal eye fields (rFEF+) is functionally necessary for endogenous [55] but not exogenous attention [56]. This double dissociation between brain regions and distinct covert spatial attention systems suggests that the neural substrates underlying related cognitive processes differ in their functional necessity. A dissociation could also exist between temporal attention and expectation. Here, we investigated whether temporal attention relies on early visual cortex: if so, disrupting V1/V2 with TMS should impair attentional behavioral benefits; if V1/V2 is not functionally necessary, these behavioral benefits should remain intact despite TMS disruption.

## Results

**Figure 1A** illustrates the experimental protocol. Observers performed a two-alternative forced-choice (2-AFC) orientation discrimination task. Throughout the experiment, observers maintained central fixation while two Gabor patches (3° in diameter, 80% contrast) were presented sequentially in the lower-right quadrant of the screen. In valid trials, an auditory precue indicated which target to attend to (T1 or T2), and the response cue matched the precue (same tone, 100% validity), instructing observers to report whether the tilt of the cued target was clockwise or counterclockwise relative to either a vertical or horizontal reference axis. In neutral trials, the precue was uninformative regarding target relevance, and the response cue instructed observers to report the orientation of either target with equal probability (50% each).

**Figure 1.**
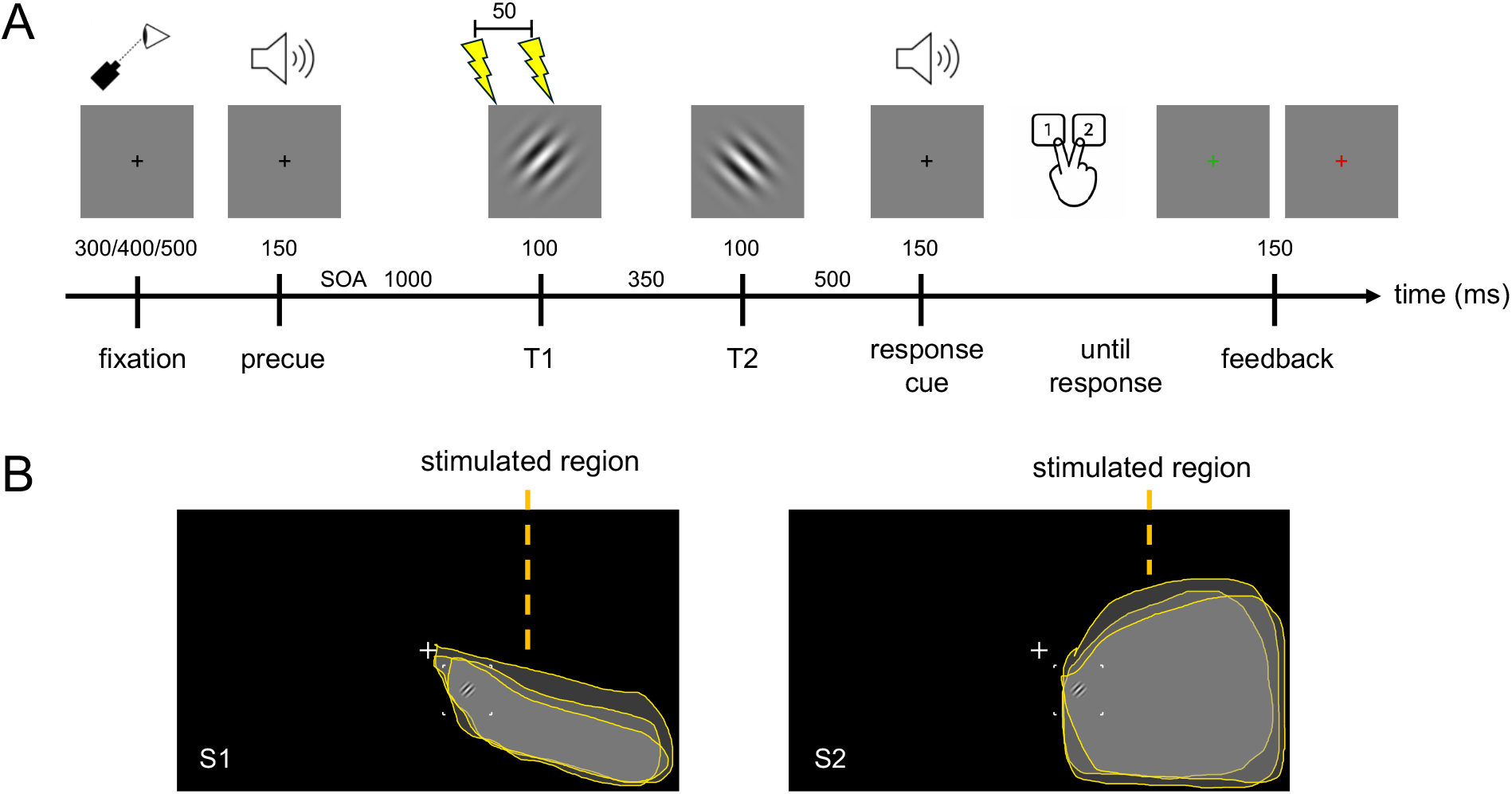
Experimental protocol. (A) Psychophysical procedure. The precue was either neutral or indicated which target’s orientation (T1 or T2) observers would be asked to report at the end of the trial (via the response cue). Observers had unlimited time to respond (with 1 indicating counterclockwise and 2 indicating clockwise) and received feedback on response accuracy (green fixation: correct; red fixation: incorrect). The lightning symbols indicate TMS pulses (50-ms interval), shown here at T1 onset as an example. (B) Phosphene mapping. Observers were stimulated near their left occipital pole and drew the outline of their perceived phosphene on the screen using a mouse. The stimulated region was tailored for each observer so that the phosphene would cover the target location in the lower-right visual field. The anatomical coordinates corresponding to this phosphene were tested at the start of each session. S1 and S2 are two representative observers, and the translucent shapes depict their drawings across all three experimental sessions. Two observers could not perceive phosphene covering the lower-right location; their targets were placed at the horizontal meridian or upper-right visual field at the same eccentricity as the other observers. Fixation marks and target spatial frequencies are not to scale.

There were four TMS conditions: (1) T1-TMS, (2) T2-TMS, (3) both-TMS (T1 and T2), and (4) none-TMS. These conditions were designed to examine whether TMS disrupts temporal attention effects on T1 and T2 while keeping expectation constant and controlling for the anticipation of the stimulation. Specifically, observers could not predict whether or when stimulation would occur, as TMS could be delivered during either target, both targets, or neither. TMS pulses were delivered at a subthreshold level using an MCF-B70 coil placed over the left occipital cortex. During phosphene mapping, we ensured that the induced phosphene covered the target location (**Figure 1B**; see Methods).

### Behavioral results

Performance was quantified using signal detection theory (see Methods). For *d′* (**Figure 2A**), a repeated-measures ANOVA (rmANOVA) with the factors of Attention (valid, neutral), Target (T1, T2) and TMS Condition (T1, T2, both, none) revealed a higher *d′* in the valid than in the neutral condition [*F*(1,10)=6.74, *p*=.027, *η*_*p*_^*2*^=.40], but no main effect of Target [*F*(1,10)<1, BF_10_=0.26] or TMS condition [*F*(3,30)<1, BF_10_=0.05], nor any interactions [3-way: *F*(3,30)=1.29, *p*=.296, BF_10_=0.21; Attention x Target: *F*(1,10)<1, BF_10_=0.272; Target × TMS: *F*(3,30)<1, BF_10_=0.08; Attention x TMS: *F*(3,30)<1, BF_10_=0.09;], indicating moderate evidence for H_0_ in the 3-way interaction and strong evidence for H_0_ in the critical Attention × TMS interaction]. Additionally, there were no significant correlations between TMS intensity and the attentional effect (valid *d′* – neutral *d′*) across observers [*ps*>.1].

**Figure 2.**
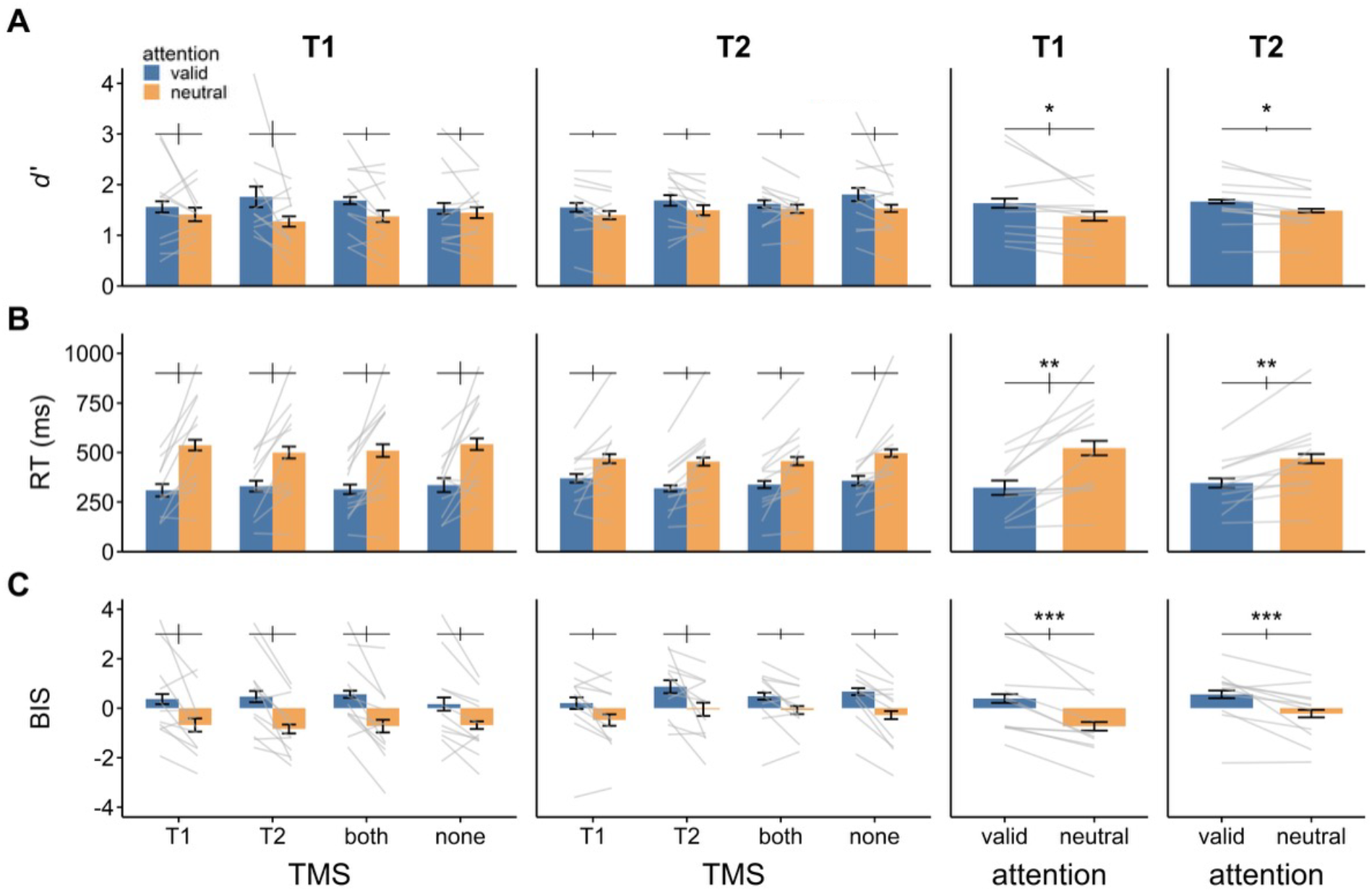
Behavioral results of the TMS-psychophysical task when T1 (left panels) and T2 (right panels) served as targets. Compared to neutral trials, valid trials yielded better performance in **(A)** *d′*, **(B)** geometric mean RT, and **(C)** BIS across TMS conditions for both T1 and T2. Gray lines indicate individual observer means. Error bars inside the bars depict ±1 SEM for each condition (Cousineau-corrected); error bars above the bars indicate ±1 SEM of the difference between conditions. *** *p*<.001, ** *p*<.01, *, *p*<.05.

For RT (**Figure 2B**), our secondary performance index, an rmANOVA on the same factors revealed faster RTs in the valid than in the neutral condition [*F*(1,10)=15.9, *p*=.003, *η*_*p*_^*2*^=.61], a main effect of TMS [*F*(3,30)=4.12, *p*=.015, *η*_*p*_^*2*^=.29], and an interaction between Attention and Target [*F*(1,10)=8.11, *p*=.017, *η*_*p*_^*2*^=.45]. No other significant differences emerged [Attention x TMS: *F*(3,30)<1, BF_10_=0.07; Target x TMS: *F*(3,30)=2.05, *p*=.128, BF_10_=0.07; 3-way: *F*(3,30)=2.27, *p*=.101, BF_10_=0.18]. Post-hoc analysis for the main effect of TMS condition revealed a faster RT in T2-TMS compared to T1-TMS [*t*(10)=2.93, *p*=.015, *d*=0.88] and none-TMS [*t*(10)=3.35, *p*=.007, *d*=1.01], but no other significant differences. Post-hoc analyses for the Attention x Target interaction reveal that the attentional benefit (RT difference between valid and neutral trials) was greater for T1 than for T2 [*t*(10)=2.85, *p*=.017, *d*=0.86].

To ensure that our findings remained consistent when accounting for both accuracy and reaction time, we conducted an rmANOVA on the Balanced Integration Score (BIS), calculated by subtracting standardized RT from standardized accuracy, with higher scores indicating better performance (**Figure 2C**). Consistent with the *d′* and RT analyses, a main effect of Attention emerged [*F*(1,10)=25.49, *p*<.001, *η*_*p*_^*2*^=.67]. No other significant effects were observed [Target: *F*(1,10)<1, BF_10_=2.86; TMS: *F*(3,30)=2.01, *p*=.134, BF_10_=0.27; Attention x Target: *F*(1,10)=3.36, *p*=.097, BF_10_=0.43; Attention x TMS: *F*(3,30)<1, BF_10_=0.07; Target x TMS: *F*(3,30)<1, BF_10_=0.09; 3-way: *F*(3,30)=1.4, *p*=.263, BF_10_=0.18].

### Pupil analysis

Pupil size was analyzed to confirm that TMS was delivered to V1/V2 at an intensity sufficient to induce physiological modulations, as pupil dilation can serve as an objective index of cortical excitability [57]. Occipital-TMS induces pupil dilation [58, 59], likely mediated by sympathetic autonomic activity, and early visual cortex activity correlates negatively with pupil size [60-62]. This is contrary to TMS applied to other brain regions, such as frontal eye fields [63, 64], or intraparietal sulcus [65], which do not induce pupil size changes.

Pupil amplitude was analyzed using the Pupil Response Estimation Toolbox (PRET) to model (**Figure 3**, see Methods) and isolate the contribution of sequential trial events [66]. We derived amplitude parameters based on model estimates. The mean *R*^*2*^ of the model estimate was 94.59%, demonstrating a strong goodness of fit. We further compared the median bootstrapped estimate with the model output; their difference [median bootstrapped estimate – model estimate] did not significantly differ from zero [*t*(9)=0.57, *p*=.581; Wilcoxon signed rank test: *V*=26, *p*=.919]. Importantly, although some observers had less eye-tracking data available, their model-fitting results were comparable to those observers with full runs.

**Figure 3.**
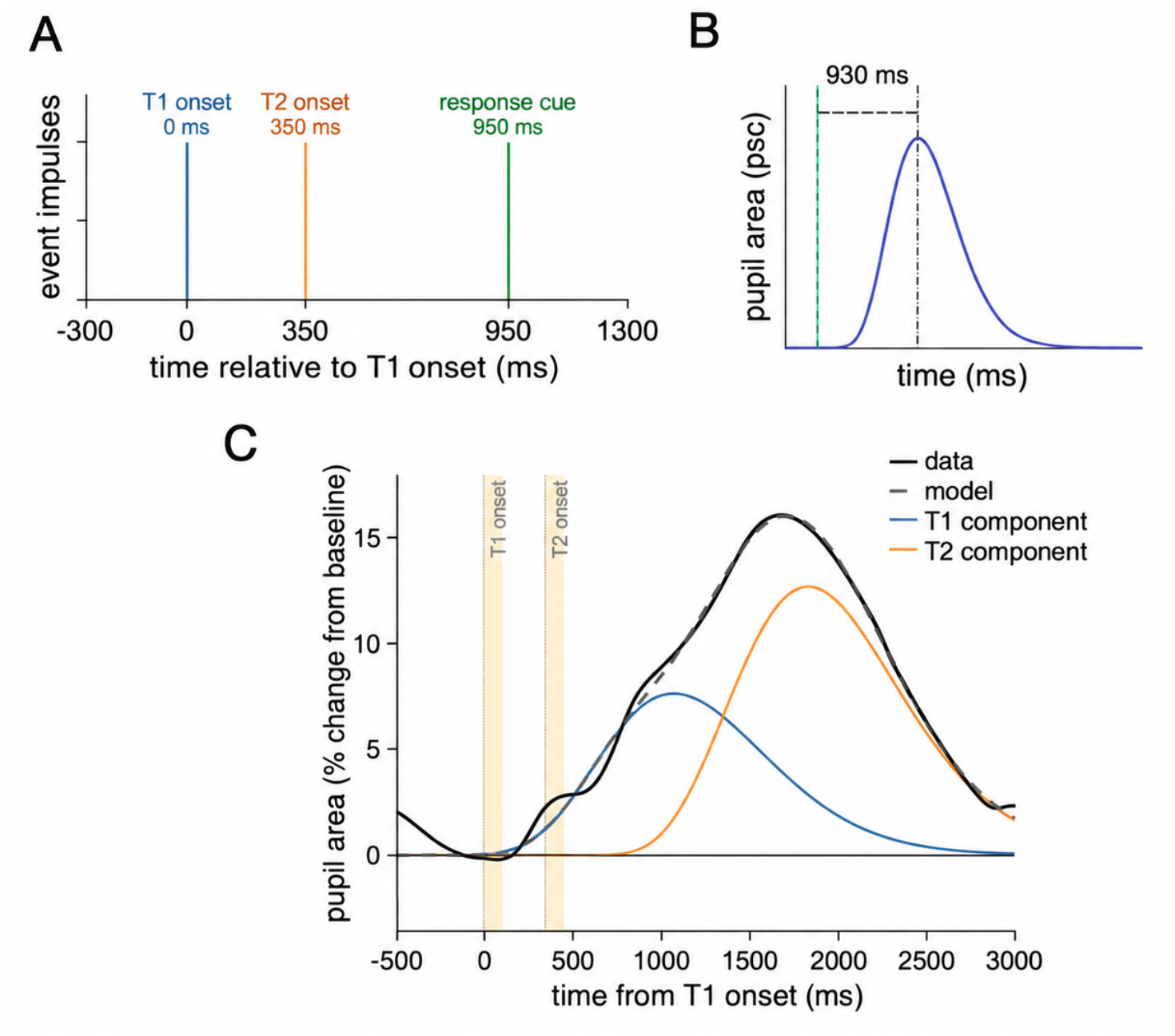
Model of the pupil response. **(A)** The modeled neural events correspond to (1) T1 onset, (2) T2 onset, and (3) response cue onset (green). Event amplitudes and latency shifts were estimated for each event **(B)** Neural event (green) and pupillary response function (Erlang gamma function, blue). The initial/default puRF peak latency was set to 930 ms, and the peak timing parameter, t_max_, was allowed to vary between 500 and 1500 ms during model fitting. **(C)** Empirical data (black) and model fit (dashed gray line) resulting from the estimated effects for each event (blue and orange). The T1 and T2 onsets are marked with yellow (100 ms).

Figure 4. shows the results for pupil dilation. After confirming that the model well captured the variance of pupil size changes, we conducted a 3-way ANOVA on the PRET-estimated event amplitude with the factors of Attention (valid, neutral), Target (T1, T2), and TMS Condition (T1, T2, both, none). A main effect of Target showed that T2 elicited a larger amplitude change than T1 [*F*(1,9)=24.52, *p*<.001]. Crucially, a main effect of TMS Condition emerged [*F*(3,27)=12.48, *p*<.001], where the both-TMS condition yielded a larger amplitude change than T1-TMS [*t*(9)=4.32, *p*=.002] and none-TMS [*t*(9)=5.39, *p*<.001]. Additionally, none-TMS yielded a smaller amplitude change than T1-TMS [*t*(9)=2.98, *p*=.016] and T2-TMS [*t*(9)=3.64, *p*=.005]. No other main effects or interactions emerged [*ps*>.1].

## Discussion

In this study, we investigated whether the early visual cortex is functionally necessary for temporal attention by applying online TMS during a two-target temporal attention task. Our experiment yielded three main results: (1) a behavioral temporal attention benefit characterized by enhanced sensitivity (*d’*), faster reaction times (RTs), and higher Balanced Integration Scores (BIS) for attended targets across four TMS conditions, (2) an absence of TMS modulation of temporal attention in the early visual cortex, and (3) significant pupil amplitude changes triggered by TMS pulses, validating the neurophysiological efficacy of our stimulation protocol. Taken together, , these results provide causal evidence that early visual cortex (V1/V2), at the time when TMS was delivered, is not functionally necessary for the voluntary allocation of temporal attention, suggesting that the causal loci of temporal attention reside outside the early visual sensory processing stream.

**Figure 4.**
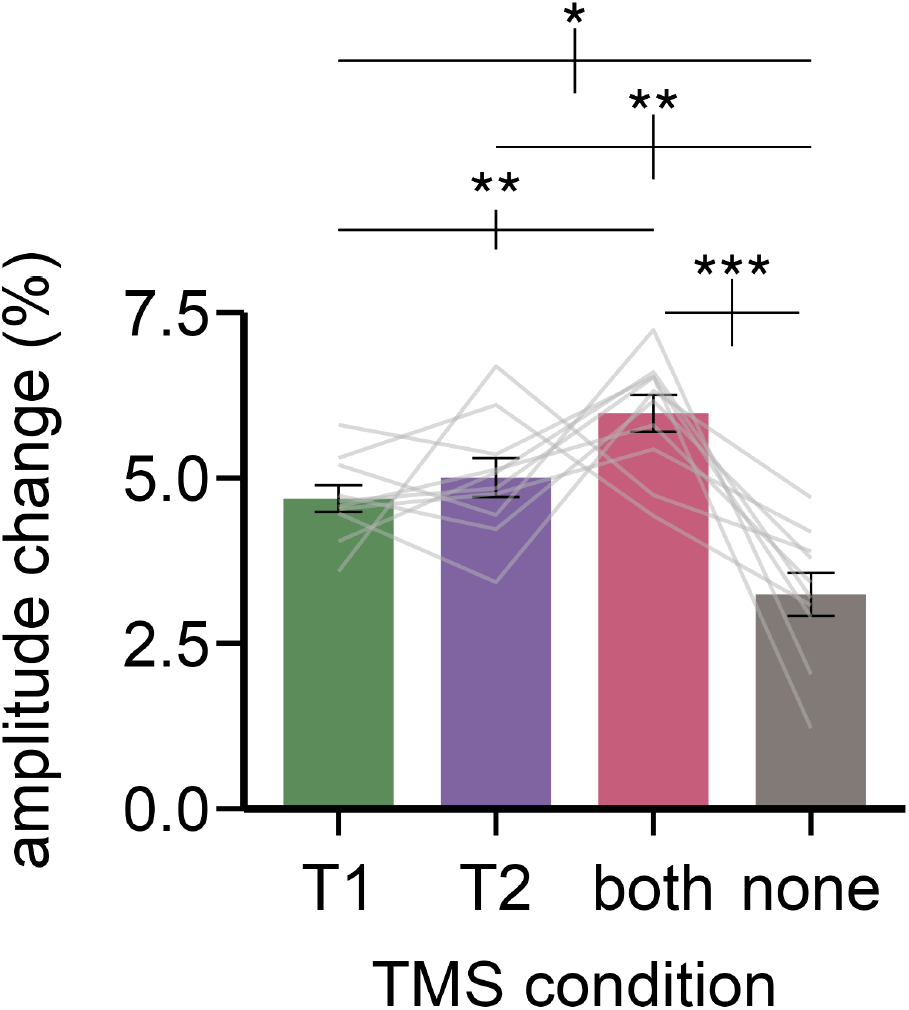
Pupil amplitude change (%) for different TMS conditions. Error bars inside the bars depict ±1 SEM for each condition (Cousineau-corrected). The gray lines indicate the mean estimates from the model across attention and target conditions for each observer in each TMS condition. Error bars above the bars indicate ±1 SEM of the difference between conditions. *** *p*<.001, ** *p*<.01, * *p*<.05.

Behaviorally, we observed typical temporal attentional performance benefits (**Figure 2**), consistent with previous studies [6-10, 67, 68]. Observers exhibited superior perceptual sensitivity (*d′*), faster RTs, and higher overall efficiency (BIS) in the validly cued condition than in the neutral condition. Crucially, for perceptual sensitivity (*d′*), there was no three-way interaction among attentional condition, target, and TMS condition, indicating that temporal attentional benefits remained consistent across targets (T1 and T2) and all four TMS conditions. For RTs, the larger attentional benefit for T1 than T2 aligned with studies showing stronger effects for earlier targets in rapid sequential streams [6, 8, 37, 38, 67], but did not interact with TMS conditions. Finally, for BIS, T2 had an overall higher BIS value than T1, driven by a lower (more negative) BIS in the neutral condition for T2 than for T1 (**Figure 2C**). Importantly, neither target asymmetry nor attentional benefits interacted with TMS conditions, confirming that disruption of early visual cortical activity did not disrupt top-down temporal attentional prioritization.

Disentangling temporal attention—voluntary information prioritization at a specific moment based on relevance—from temporal expectation—predicting when an event will occur based on temporal structure has proven challenging, leading many previous studies to conflate these constructs. Although temporal attention and expectation frequently co-occur in ecological settings and share overlapping functional goals [17], they may rely on distinct neural mechanisms. For instance, temporal expectation enhances visually evoked responses [27, 28]. In contrast, voluntary temporal attention improves task performance without necessarily altering early sensory-evoked responses in primary visual cortex [37, 38]. Indeed, when both are independently manipulated, they interact: attentional benefits decrease with delayed target onset despite increased expectation [8]. We kept temporal expectation constant here; thus, our findings are agnostic regarding temporal prediction, but they provide causal evidence that endogenous temporal attention does not require early visual cortex. By demonstrating that disrupting V1/V2 leaves temporal attention intact, our results reinforce the theoretical dissociation between temporal expectation and voluntary temporal attention [4, 6, 7], supporting the view that temporal attention relies on higher-order brain regions [13, 37, 38, 69].

If early visual areas V1/V2 are not functionally required for temporal attention, what candidate brain structures mediate these selective perceptual gains? Recent MEG evidence implicates left fronto-cingulate cortex, showing elevated target orientation-decoding accuracy during attended than unattended conditions, specifically for T1 [38]. Furthermore, a recent study reports that endogenous temporal attention dynamically routes T1 target sensory representations to fronto-cingulate cortex, which may protect them from backward masking and temporal interference when T2 subsequently appears [39]. The functional locus of temporal attention may thus reside within higher-order networks responsible for maintaining, shielding, and prioritizing task-relevant representations against competitive sensory input. Consequently, disruption of V1/V2 via TMS fails to eliminate the attentional advantage, as the critical selection and gating operations likely take place downstream in frontocortical regions beyond the early visual pathway.

We tested four TMS conditions in the current study to examine whether the temporal attention effect on T1 and T2 would be selectively diminished while controlling the behavioral expectation effects of TMS pulses [70]. Interleaving these four TMS conditions (T1-TMS, T2-TMS, both-TMS, and none-TMS) across trials, ensured that observers could not anticipate whether, when, or how many pulses would occur. Furthermore, T1-TMS and T2-TMS served as mutual control conditions: T1-TMS should not affect T2 processing, and T2-TMS should not affect T1 processing, given that occipital TMS effects last at most ∼140 ms [71] shorter than the 350-ms T1-T2 stimulus onset asynchrony (SOA). Crucially, pulses at T1 provided no predictive information regarding stimulation at T2. Therefore, this design ensured that the lack of TMS modulation was not driven by pulse anticipation.

When interpreting the lack of effect of TMS on V1/V2, it is important to keep in mind that the stimulation protocol is capable of eliminating the effects of attention; the same stimulation protocol eliminated the effects of exogenous spatial attention in a very similar task [53, 54] and disrupted visual working memory maintenance in a memory-guided saccade task [72]. In general, when interpreting null behavioral effects in brain stimulation research, it is important to ensure that the delivered stimulation was sufficient to induce neural disruption [73, 74]. Multiple lines of physiological and computational evidence confirm that our TMS protocol achieved robust efficacy.

First, analyzing pupil dynamics using the Pupil Response Estimation Toolbox (PRET) [66]—revealed clear physiological responses to stimulation. Although pupil dilation did not vary across attentional conditions —aligning in both pattern and magnitude with prior work [66]—it showed strong sensitivity to TMS stimulation: dilation was largest in the both-TMS condition, followed by T1-TMS and T2-TMS, all exceeding the none-TMS condition (**Figure 4**).

This finding aligns with previous occipital TMS studies using different TMS protocols and repetitive stimulation intensity of 110% or 130% of motor threshold [58, 59]. Pupil dilation has also been reported following peripheral stimulation of the occipital nerves in humans [75], stimulation of the visual cortex in other mammalian species [76, 77], and optogenetic stimulation of locus coeruleus axons within the primary visual cortex in mice [78]. Although we cannot fully exclude a contribution from the acoustic effects of TMS, evidence that active TMS elicits greater pupil dilation than sham stimulation designed to preserve the coil click [57] suggests that auditory stimulation alone is unlikely to fully account for the pupil dilation observed here.

Second, individual thresholding procedures confirmed high stimulation potency. Consistent with previous studies [79-81], individual phosphene thresholds in our sample exceeded motor thresholds, with the applied stimulation intensity averaging 1.39 times the individual motor thresholds (**Table S1**). This stimulation matches or exceeds the relative intensities established to induce pupil dilation in visual cortex studies [58, 59].

Third, biophysical electric field (E-field) simulations (SimNIBS ver. 4.6.0 [82]) yielded an average induced peak cortical E-field of 1.13 V/m (53.27% MSO, **Figure S1**). This induced field strength is comparable to levels that successfully eliminated exogenous spatial attention effects in V1/V2 using a different TMS system (Magstim Rapid Plus stimulator 3.5T; 1.32 V/m in [53]; 1.31 V/m in [54]), and a higher average stimulation intensity disrupted visual working memory maintenance in early visual cortex using the same TMS system (49.28% MSO; [72]). Thus, given the pupil dilation results, rigorous localization, and stimulation intensities exceeding established functional thresholds [73, 74], our stimulation protocol was physiologically effective and appropriately targeted. The absence of behavioral modulation therefore reflects functional independence of endogenous temporal attention from early visual cortical processing rather than an underdosed or otherwise ineffective stimulation protocol.

The TMS effect on attention can differ between performance benefits and costs. For example, in presaccadic spatial attention, which enhances processing at a target location just before a saccadic oculomotor response, stimulating V1/V2 right before saccade onset reduced performance benefits without modulating costs, whereas stimulating rFEF+ significantly reduced costs while leaving benefits intact [83]. Thus, these brain regions play distinct roles in modulating presaccadic attentional benefits versus costs. Because our experimental design focused on measuring attentional benefits (valid vs. neutral cues), we cannot rule out the possibility that the early visual cortex plays a functional role in mediating the costs (neutral vs. invalid cues) of temporal attention.

Endogenous temporal attention may also rely on distinct neural underpinnings compared to exogenous temporal attention, as the former is voluntary and goal-driven, whereas the latter is involuntary and stimulus-driven [84]. These differences parallel classic distinctions between endogenous and exogenous spatial attention [2]. TMS-psychophysical studies examining spatial attention show that V1/V2 is critical for exogenous [53, 54] but not endogenous attention [55], whereas the rFEF is critical for endogenous [55] but not exogenous spatial attention [56]. This pattern raises the possibility that early visual cortex is unnecessary for endogenous processes across both spatial and temporal domains. Furthermore, because existing evidence linking the fronto-cingulate network to endogenous temporal attention is correlational [37, 38, 69], future brain stimulation studies should directly test the causal involvement of these higher-order regions.

A previous non-invasive brain stimulation study examining the neural substrates of temporal processing has jointly manipulated temporal attention and expectation. For instance, transcranial alternating current stimulation (tACS) over the right posterior parietal cortex abolishes the temporal expectation effect under attended conditions without altering temporal attention per se [11]. Other studies, which did not isolate temporal attention and expectation, applying TMS over the intraparietal sulcus (IPS) failed to modulate performance during temporal-expectation tasks [15] or demonstrated that repetitive TMS over the right temporoparietal junction (TPJ) impaired simultaneity-judgment tasks [85]. Together, these findings suggest that different aspects of temporal information processing rely on distinct neural mechanisms. By clearly defining and manipulating voluntary temporal attention, while keeping temporal expectation constant, our study fills a critical gap in the literature, showing that voluntary temporal attention operates through a distinct architecture for which early visual sensory nodes are not necessary.

In conclusion, our findings demonstrate that early visual cortex is not functionally necessary for voluntary temporal attention. These results help dissociate endogenous temporal attention from temporal expectation, aligning instead with studies implicating higher-order fronto-cingulate regions in supporting temporal attention by preserving and shielding task-relevant sensory representations over time.

## Methods

### Participants

To ensure sufficient power to detect both temporal attention and TMS effects on attention, we conducted power analyses using G*Power 3.1 [86] based on effect sizes reported by Denison, Heeger and Carrasco [6] for temporal attention (*d*=1.21) and Lee, Fernández and Carrasco [54] for the Attention × TMS interaction (*η*_*p*_^*2*^=.26). The analyses indicated that 10 observers were required to detect the temporal attention effect, and 7 observers were required to detect the Attention × TMS interaction (α=.05, 90% power, assuming a correlation among repeated measures of *r*=.50 and a nonsphericity correction of *ϵ*=1). Therefore, eleven observers (5 females; age range: 22-42, mean age: 28.55), including author H.H.L., participated in the experiment. This sample size is equivalent to or greater than the typical number of observers in previous perceptual and psychophysical TMS studies [e.g., 52-54, 83, 87, 88-92]. All observers (except the author) were naïve to the purpose of the study and provided written informed consent prior to participation in the experiment. All observers had normal or corrected-to-normal vision and no history of neurological disorders. The study adhered to the safety guidelines for TMS research and was approved by the New York University Langone Health Institutional Review Board (IRB protocol i14-00788_MOD20).

### Apparatus

Stimuli were presented on a gamma-calibrated ViewPixx LCD monitor (120 Hz refresh rate, 1920 × 1080 resolution). An EyeLink 1000 eye tracker (SR research) recorded observers’ gaze (right-eye position, with 1,000 Hz sampling rate) to ensure continuous central fixation throughout each trial and confirm that attention was covert. If observers moved their eyes (deviation > 1.5 dva), the trial was interrupted and repeated at the end of the block.

### Stimuli

Stimuli were displayed on a uniform medium gray background (72.92 cd/m^2^). A central fixation cross (0.25° × 0.25°, 0.06° thickness) was present throughout each trial. Four small black placeholders (0.28° × 0.28°, 0.06° thickness) placed at the corners of an imaginary square (5.28° × 5.28°) centered in the lower right of the visual field at 6° eccentricity (following [6]). For the two observers who could not perceive phosphenes at this location, placeholders and targets were relocated to the horizontal meridian and upper-right visual field, respectively, at the same 6° eccentricity. Targets were placed in the right visual field based on evidence that temporal attention is left-lateralized [5, 13, 38, 39].

Target Gabors (diameter = 3°, spatial frequency = 3 cycle/degree (cpd), full width at half maximum = 1°) were presented at 80% contrast with a random phase. This contrast level was chosen because temporal attention typically increases response gain on the contrast response function [68]. Each target was tilted clockwise (CW) or counterclockwise (CCW) relative to either a vertical or horizontal reference axis. Tilt angles were titrated independently for each observer, target (T1 vs. T2) and reference axis.

Auditory stimuli were delivered via external speakers. The 150-ms auditory precue was either a pure sinusoidal tone or a complex waveform composed of sinusoidal frequencies from 50 to 400 Hz. A high-frequency tone (1,000 Hz) cued the first target (T1), a low-frequency tone (300 Hz) cued the second target (T2), and the complex tone served as an uninformative neutral precue. The same T1 or T2 tones were also presented as response cues at the end of the trial to instruct the observer to report the orientation of T1 or T2, respectively.

### TMS and phosphene mapping

TMS pulses were delivered using an MCF-B70 coil placed over the left occipital cortex, connected to a MagVenture MagPro X100 stimulator and triggered via a MATLAB-controlled Arduino board. Stimulation intensity remained constant across experimental sessions for each observer and was set according to their individual phosphene threshold, where the psychophysical task was performed with the screen presenting a gray background and the phosphene mapping was performed with the screen in a complete black background (40%–59% of maximum stimulator output, mean = 53.27%, SD = 5.22%). The phosphene mapping protocol followed established procedures same as in previous studies [53-55, 72, 88-90, 93].

Observers were seated 57 cm from the monitor in a dark room and fixated on a dark-blue cross on a black background. A train of seven TMS pulses (30 Hz at 60% maximum stimulator output) was applied to the left occipital cortex. Observers drew the outline of the perceived phosphene on the screen using a mouse, and the coil position was adjusted so that the perceived phosphene overlapped with the target location in the lower-right visual field, except for two observers who could not perceive the phosphene in the lower-right visual field, their targets were placed at horizontal meridian and upper-right visual field, where they could perceive phosphenes consistently, with the same eccentricity as other observers.

Phosphene thresholds were determined using two pulses spaced 50 ms apart at the identified coil location [53-56, 83, 88, 89, 93, 94]. Pulse intensity was adjusted until observers reported perceiving phosphenes on 50% of the trials. In addition to phosphene thresholds, we measured observers’ motor thresholds by stimulating the right motor cortex and monitoring visible twitches of the left hand. Motor threshold was defined as the pulse intensity that elicited a visible twitch on 50% of trials. Each observer’s head was coregistered to a 3D template in Brainsight Neuronavigation software, to ensure millimeter-level precision across trials. Coil coordinates were saved in Brainsight Neuronavigation software. The stimulation intensity, final stimulation MNI coordinate, and motor threshold information are listed in **Table S1**.

### Experimental procedure

Observers performed a two-alternative forced-choice (2-AFC) orientation discrimination task. Throughout the experiment, observers maintained central fixation while two targets were presented sequentially at the lower right quadrant of the screen. For two observers who could not perceive phosphene in the lower right visual field, targets were placed at the same eccentricity along the horizontal meridian and in the upper right quadrant, respectively.

**Figure 1A** illustrates the experimental protocol. Each trial began with a central fixation cross presented for 300, 400, or 500 ms, followed by a 150-ms auditory precue indicating whether to attend to the first target (T1), second target (T2) or both targets. Following the precue, a 1000-ms delay before each target was presented for 100 ms with a stimulus onset asynchrony (SOA) of 350 ms. This SOA was selected because temporal attention induces the strongest attentional effects at SOAs between 200-450 ms [7] and occipital-TMS effects typically persist for less than 140 ms [71]. An auditory 150-ms response cue was presented 500 ms after the offset of the second target. In valid trials, the precue indicated which target to attend (T1 or T2), and the response cue was the same tone as the precue (i.e., 100% validity), instructing observers to report the orientation of the cued target. In neutral trials, the precue was uninformative regarding target relevance, and the response cue instructed observers to report the orientation of either target with the same probability (50%).

The Gabor stimuli had one of two reference orientations: vertical or horizontal. T1 and T2 could share the same orientation or differ, as in previous temporal attention studies, e.g., [4, 6-9, 67, 68, 84]. Observers were required to determine the relevant reference orientation on each trial. Feedback was presented for 300 ms following each response: the fixation cross turned green for a correct response and red for an incorrect response. This was followed by a 500-ms intertrial interval (ITI). Observers had no time limit to respond and were instructed to prioritize accuracy over speed. Eye movements were monitored continuously throughout the experiment, and observers were instructed to maintain central fixation.

There were four TMS conditions, (1) T1-TMS, (2) T2-TMS, (3) both-TMS (T1 and T2), and (4) none-TMS. These conditions were designed to examine whether TMS would disrupt temporal attention effects on T1 and T2 while controlling for expectation effects. Specifically, observers could not predict whether and when stimulation would occur, as TMS could be delivered during either target, both targets, or neither.

The experiment included two attentional conditions (valid and neutral), two target conditions (T1 and T2), and four TMS conditions (T1, T2, both, and none TMS). Each block contained 64 trials. Each observer completed eight blocks per session across three sessions, for a total of 1,536 trials.

Before beginning the TMS-psychophysical task, observers completed a titration task to estimate tilt-angle thresholds for T1 and T2 for each vertical and horizontal reference orientation. Two staircases were conducted for each combination of target and reference orientation: one beginning at 0.5° and the other at 7.5°. Tilt-angle thresholds were estimated at 79% accuracy using an adaptive staircase procedure implemented with the Palamedes toolbox [95], and following established methods [53, 54].

For each observer, tilt-angle thresholds for each target and reference orientation were calculated by averaging the values from the final 5 trials of two staircases. Each staircase contained 30 trials for each of the four possible staircase combinations (horizontal-reference T1, horizontal-reference T2, vertical-reference T1, and vertical-reference T2), yielding 60 trials per block and 240 trials across four blocks.

### Statistical analysis for behavioral data

Sensitivity was indexed by *d′* [z(hit rate) – z(false alarm rate)] across conditions. Correct discriminations of clockwise trials were classified as “hits” and incorrect discriminations of counterclockwise trials were classified as “false alarms”, as in previous studies, e.g., [96-99].

When analyzing reaction time (RT) data as a secondary performance index, we included only correct trials and computed geometric mean RTs to minimize the influence of unusually long responses within the typically right-skewed RT distribution [100], following previous studies, e.g., [101-104].

We also accounted for both accuracy and speed simultaneously by calculating the Balanced Integration Score (BIS) [105]. The BIS assigns equal weight to accuracy and RT by converting both into z-scores, placing them on the same scale and subtracting the RT z-score from the percent correct z-score. BIS is commonly used in visual attention studies, e.g., [8, 9, 106-108], to ensure that findings are not confounded by speed-accuracy tradeoffs.

Bayes factors were estimated using the BayesFactor package in R [109] to evaluate evidence for the null hypotheses. Default Jeffreys–Zellner–Siow (JZS) priors were used, with fixed effects assigned a medium scale (*r*=0.5) and random effects a nuisance scale (*r*=1). BF_10_ values favoring the null hypothesis were categorized as anecdotal (0.33–1), moderate (0.10–0.33), strong (0.033–0.10), very strong (0.01–0.033), or extreme (<0.01). Conversely, evidence favoring the alternative hypothesis was categorized as anecdotal (1–3), moderate (3–10), strong (10–30), very strong (30–100), or extreme (>100) [110].

### Pupil data preprocessing and modeling

Pupil area size was analyzed to confirm that TMS was delivered to V1/V2 at an intensity sufficient to induce physiological modulations. Occipital-TMS induces pupil dilation [58, 59], likely mediated by sympathetic autonomic activity, and visual cortex activity correlates negatively with pupil size [60-62].

Pupil data from 10 of the 11 observers were analyzed; data from one observer were excluded due to a technical recording error. Of the 10 observers included in the analysis, six had complete data from all three runs, three had data from two runs, and one had data from a single run.

Raw pupil size data were epoched into trial-wise segments. Due to the long, sluggish kinetics of pupillary responses, blinks were unlikely to affect low-frequency changes in pupil amplitude [111, 112]; thus, to produce a smooth time series for model fitting, blink periods were reconstructed using cubic-spline interpolation [112]. Specifically, during the experiment, fixation was monitored online during the pre-cue/target period. Trials with fixation failures during these ‘checkpoints’ were aborted and re-tested at the end of each block. The response-cue period was not used for online trial abortion. Offline pupil preprocessing was therefore applied to completed trials and primarily handled blink periods or brief pupil losses that remained in the epoch, including blinks after the online monitoring window. EyeLink-defined blink periods were extended by 100 ms before blink onset and 100 ms after blink offset; samples within these padded windows, as well as zero, negative, or missing pupil samples, were marked as invalid and linearly interpolated within each epoch. Pupil time series were converted to percent signal change from a –200 to 0 ms relative to T1 onset, pre-stimulus baseline, using PRET to model preprocessing.

Pupil time series were modeled using a general linear model (GLM) framework [66]. Pupil size was estimated on a trial-wise basis, as the sum of dilatory responses for two neural events (impulses) convolved with a pupil response function (puRF). Each impulse had free parameters for amplitude (% signal change) and peak (ms), representing the internal signals associated with trial event onsets. Convolved impulses were fitted to pupil responses evoked by T1 and T2 using T1-locked epochs from –500 to 3,500 ms relative to T1 onset. The model included two impulse events corresponding to T1 onset (0 ms) and T2 onset (350 ms; **Figure 3A**). Event amplitudes were constrained between –10% and 30% signal change from baseline. Event impulses were convolved with the PRET pupil response function, modeled as an Erlang gamma function. The default t_max_ value was set to 930 ms (**Figure 3B**) but was allowed to vary between 500 and 1,500 ms. No sustained boxcar regressor, response-cue regressor, intercept, or linear slope term was included in the model.

For PRET fitting, trials with more than 30% blink-marked samples or fewer than 70% finite pupil samples within the epoch were excluded (23 out of 12,352 trials, exclusion rate: 0.19%). Incorrect trials were retained. Model fitting was performed on the condition-mean pupil time course for each observer and condition following trial exclusion, with runs pooled within observer and condition. For each observer × condition, the five estimated free parameters were two event amplitudes, two event latency shifts, and the t_max_ parameter. Parameter estimation used 2,000 sampled parameter sets, with the best 40 serving as starting points for constrained optimization. Together, these reconstructed signals were summed to predict pupil dynamics across each trial segment (**Figure 3C**). To ensure that observers with fewer available runs yielded stable model estimates, we conducted bootstrapping for each condition per observer (100 times) [66] and compared the bootstrapped median of each parameter with the corresponding model estimate. Estimates were deemed stable when the bootstrapped median closely matched the original model estimate.

## Acknowledgments

The authors thank Nina Hanning, Caroline Myers, and Nick Crotty, and other members of the Carrasco Lab, as well as Rachel Denison, for their feedback on a previous version of the manuscript.

