## Supplementary Materials for "Early visual cortex is not functionally necessary for temporal attention: A TMS study"

### Table S1

#### Stimulation loci, stimulation intensity, and motor thresholds

The stimulation loci are individualized through the phosphene induction procedure at the beginning of each session. The phosphenes were short lasting spatially circumscribed percepts reported in the hemifield contralateral to the stimulated occipital pole. The stimulating loci where consistent throughout the experiment while maintaining the error being smaller than 1 mm.

| observer number | stimulation intensity (MSO%) | E-field (V/m) | Target MNI coordinate [x,y,z] | motor threshold (MSO%) |
|---|---|---|---|---|
| 1 | 54 | 1.11 | -38.23, -132.91, 71.86 | 37 |
| 2 | 54 | 1.02 | -44.85, -113.29, 76.66 | 43 |
| 3 | 56 | 1.25 | -38.34, -132.33, 26.3 | 37 |
| 4 | 56 | 1.22 | -21.62, -119.63, 45.91 | 44 |
| 5 | 52 | 1.17 | -39.29, -123.47, 62.82 | 38 |
| 6 | 49 | 0.93 | -48.5, -106.8, 92.84 | 31 |
| 7 | 40 | 1.06 | -51.78, -127.92, 81.35 | 36 |
| 8 | 53 | 1.12 | -34.55, -121.19, 69.58 | 38 |
| 9 | 59 | 1.10 | -74.69, -110.46, 54.78 | 41 |
| 10 | 58 | 1.24 | -36.56, -132.65, 46.83 | 40 |
| 11 | 55 | 1.18 | -46.86, -128.95, 39.98 | 35 |

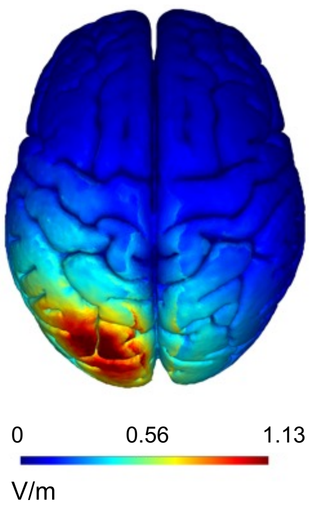


**Figure S1.** Simulated cortical excitability. Estimated E-field for V1/V2 using the simNIBS toolbox.
